# Quantitative Profiling of Lysosomal pH Heterogeneity using Fluorescence Lifetime Imaging Microscopy

**DOI:** 10.1101/2023.09.25.559395

**Authors:** Dinghuan Deng, Baiping Wang, Youchen Guan, Hui Zheng, Ayse Sena Mutlu, Meng C. Wang

## Abstract

Lysosomes play crucial roles in maintaining cellular homeostasis and promoting organism fitness. The pH of lysosomes is a crucial parameter for their proper function, and it is dynamically influenced by both intracellular and environmental factors. Here, we present a method based on fluorescence lifetime imaging microscopy (FLIM) for quantitatively analyzing lysosomal pH profiles in diverse types of primary mammalian cells and in different tissues of the live organism *Caenorhabditis elegans*. This FLIM-based method exhibits high sensitivity in resolving subtle pH differences, thereby revealing the heterogeneity of the lysosomal population within a cell and between cell types. The method enables rapid measurement of lysosomal pH changes in response to various environmental stimuli. Furthermore, the FLIM measurement of pH-sensitive dyes circumvents the need for transgenic reporters and mitigates potential confounding factors associated with varying dye concentrations or excitation light intensity. This FLIM approach offers absolute quantification of lysosomal pH and highlights the significance of lysosomal pH heterogeneity and dynamics, providing a valuable tool for studying lysosomal functions and their regulation in various physiological and pathological contexts.

## INTRODUCTION

Lysosomes are dynamic organelles that function as the primary site for macromolecular catabolism. The metabolic function of lysosomes relies on their acidic pH, which is a unique feature of these specialized cellular organelles (Forgac, 2007). A variety of lysosomal hydrolases reside in this acidic compartment and degrade macromolecules into amino acids, monosaccharides, nucleotides, and free fatty acids (Perera and Zoncu, 2015). Lysosomes are functionally and morphologically heterogenous and change their activities in response to various intra-and inter-cellular inputs (Perera and Zoncu, 2015). Lysosomal functions are crucial for the maintenance of cellular homeostasis and organism fitness. Dysfunction of lysosomes underlines the pathology of multiple chronic and degenerative diseases (Colacurcio and Nixon, 2016; Peng *et al*., 2019). In particular, lysosomal storage diseases due to deficiencies in lysosomal degradation show neurological deficits related to alterations in both glial and neuronal functions (Beck, 2016; Lie and Nixon, 2018; Kreher *et al*., 2021), and lysosomal dysfunction has been implicated in the development of Alzheimer’s disease as well (Zhang *et al*., 2022). Lysosomal pH is a key indicator of lysosomal function, and quantitatively measuring lysosomal pH *in vivo* is essential for determining changes in lysosomal functions under various physiological and pathological conditions.

Several pH-sensitive fluorescent protein variants have been developed to observe the pH of intracellular organelles based on fluorescence intensity and fluorescence ratio imaging (Han and Burgess, 2010; Kim and Seong, 2021). These proteins can be targeted to lysosomes as pH indicators through genetically engineering cells, tissues, and organisms. Unfortunately, these genetic engineering approaches are not easily applicable in primary cell cultures and may require labor-intensive, time-costly procedures to generate transgenic animals. On the other hand, pH-sensitive chemical probes can be directly applied for staining lysosomes. However, the robustness of the measurement based on their fluorescence intensity can be influenced by multiple factors, such as probe absorption and distribution heterogeneity, fluorophore concentration and photobleaching, and fluctuation in excitation light. To avoid these confounding effects in the intensity measurement, ratiometric fluorescence imaging methods utilize a stable internal control and simultaneous acquisition of fluorescent signals at two different excitation and/or emission wavelengths (Burgstaller *et al*., 2019; Chin *et al*., 2021; Ponsford *et al*., 2021; Webb *et al*., 2021). The other way to override the limitation of the intensity measurement is to conduct fluorescence lifetime imaging, which measures the time a fluorophore remains in an excited state before emitting a photon. The fluorescence lifetime is affected by the chemical environment surrounding the fluorophore but independent of fluorophore concentration and excitation light intensity (Datta *et al*., 2020). One key environmental factor contributing to the change in fluorescence lifetime is pH. As a result, fluorescence lifetime imaging microscopy (FLIM) provides a quantitative way for pH measurements.

Here, we have used a pH-sensitive probe that is specifically taken up into lysosomes, and applied FLIM to monitor the lifetime change of this probe in primary mammalian cells and live animals, *Caenorhabditis elegans (C. elegans).* This allows us to rapidly, quantitatively measure lysosomal pH at organellar resolution *in vivo*. We discovered that lysosomes are more heterogeneously distributed across a larger range of pH in primary microglial cells than in primary astrocytes. We also discovered that during the maturation process of primary neurons, lysosomes become more acidic and less heterogeneous, and lysosomes in mature neurons have a lower pH than those in primary astrocytes and microglial cells. The activation of microglial cells decreases lysosomal pH. Furthermore, we discovered that serum starvation increases lysosomal pH in primary microglial cells and neurons. In live worms, we revealed the tissue specificity of lysosomal pH at the whole organism level. Together, these studies demonstrate the power of FLIM in quantifying lysosomal pH and revealing its dynamic changes over time and space in various biological systems under different environmental conditions.

## RESULTS and DISCUSSIONS

### FLIM system setup and image analysis

The FLIM system is based on an ISS Q2 laser scanning confocal nanoscope, which is equipped with one-photon excitation laser launcher, fastFLIM data acquisition and processing units (Figure 1A). The data acquisition card of the FLIM system is developed using digital frequency domain technique that allows the acquisition of time-tagged-time-resolved data without the dead time typically of the time-correlated single photon counting (TCSPC) approach. Using the “digital frequency-domain” technique, the excitation light is modulated at a frequency ω, and the phase shift φ and the modulation *m* are measured for each pixel location (*h, k*). The phase shift φ and the modulation *m* are then used to calculate the coordinate g(ω) and s(ω) that are the real and imaginary components of the Fourier Transform of the fluorescence decay function, respectively at a single exponential decay. Thus, the fluorescence lifetime of pixels can be shown on a phasor plot that is the representation of a semicircle in the (s, g) plane with the center at (0.5, 0) and the radius of 0.5 (Figure 1B). On this circle, a phasor corresponding to a short lifetime (small φ) is close to the coordinate (1,0), whereas a phasor corresponding to a long lifetime (big φ) is close to (0, 0) (Digman *et al*., 2008).

**Figure 1.**
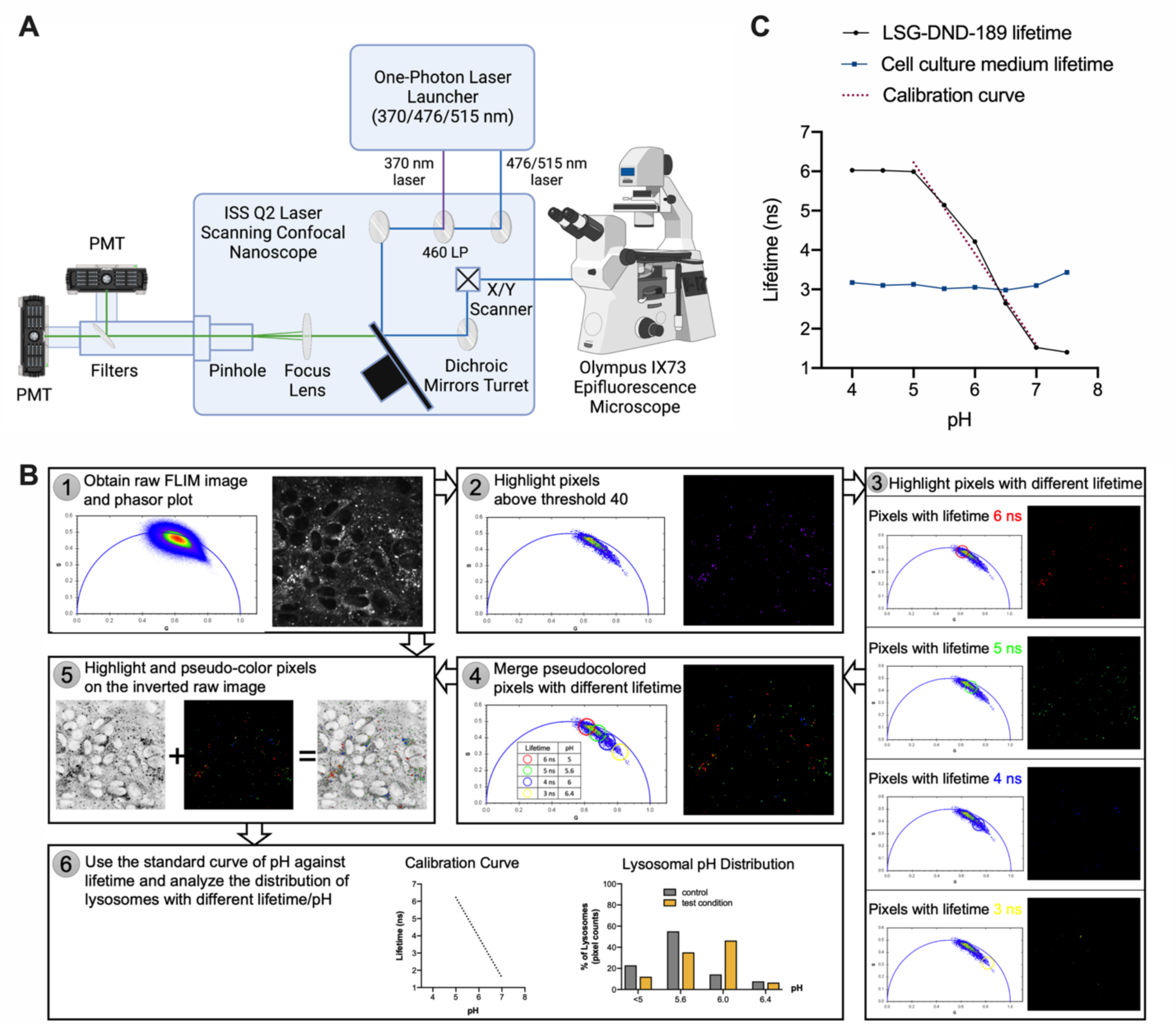
FLIM system setup and image analysis. (A) Scheme of the FLIM system. (B) FLIM image analysis pipeline showing the raw FLIM image, phasor plot, the process of highlighting pixels with different lifetime and merging pseudocolored pixels based on lifetime. (C) pH-lifetime calibration curve based on lifetime measurement of LSG-DND-189 in culture medium and lifetime analysis of culture medium alone across different pH.

LysoSensor Green DND-189 (LSG-DND-189) is a vital dye that accumulates in acidic organelles, like lysosomes. Its fluorescent intensity increases as the acidity of the lysosomes decreases and it is almost nonfluorescent in nonacidic compartments (Perzov *et al*., 2002). Previous studies demonstrated that the fluorescence lifetime of LSG-DND-189 exhibits a pH-dependent change (Lin *et al*., 2001). In our studies, we implemented fastFLIM to measure the decay time of LSG-DND-189 in frequency domain. We first established a pH calibration curve using the culture medium for growing primary cells and titrated LSG-DND-189 between pH 4.0 and 7.5 at increments of 0.5 (Figure 1C). To verify there is no intensity contribution from the culture medium, we also imaged the cell culture medium without LSG-DND-189 between pH 4.0 and 7.5 using the same imaging settings. We found that the cell culture medium has a lifetime around 3 ns at all pH conditions (Figure 1C), but the average fluorescent intensity derived from the cell culture medium alone was negligible compared to the intensity of LSG-DND-189 with the medium (Supplementary Figure 1).

We found that the calibration curve is linearly correlated with fluorescence lifetime in the range of pH 5.0 to 7.0, and thus, the linear region of the calibration curve can be fitted with the following equation with Pearson correlation coefficient, r= -0.99376:

pH = 7.68464 - 0.43152 x Lifetime

This calibration curve presented a dynamic range of lifetime changes in LSG-DND-189 between 6.0 ns and 1.5 ns that are in reverse correlation with pH 5.0 and 7.0. The LSG-DND-189 dye has a pKa of 5.2. As a result, DND-189 is unable to further distinguish pH changes below 5 (Figure 1C), and thus, those pixels (6.0 ns) if present would be assigned a pH value of <5.0 (Figure 1B).

In order to measure lysosomal pH changes *in vivo*, we processed and analyzed FLIM imaging datasets using the ISS VistaVision software and ImageJ (Figure 1B). We set an intensity threshold to select out lysosomal pixels and eliminate background pixels. We next highlighted and pseudo-colored the pixels with different lifetimes (6.0 ns, 5.0 ns, 4.0 ns, 3.0 ns correlated with pH <5.0, 5.6, 6.0 and 6.4, respectively) on the phasor plot and measured their numbers. Finally, we quantified the percentage of the pseudo-colored pixels with different lifetimes among all highlighted pixels to determine the distribution of lysosomes across different lysosomal pH levels (lysosomal pH distribution, Figure 1B).

### Lysosomal pH difference between astrocytes and microglia

Next, we applied this method in primary cell cultures from isolated cerebral cortices of P3 newborn mice and examined whether lysosomal pH exhibits any difference between glial cells. Astrocytes were obtained directly by culturing the cell suspension; while for microglia, additional steps were included with a final transfer of floating cells. The preparation of primary glial cells after isolating the brain from mice typically took 7-10 days. Then, both glial cell types were stained with LSG-DND-189 for 5 minutes, washed with fresh culture medium, and used for FLIM imaging (Figure 2A).

**Figure 2.**
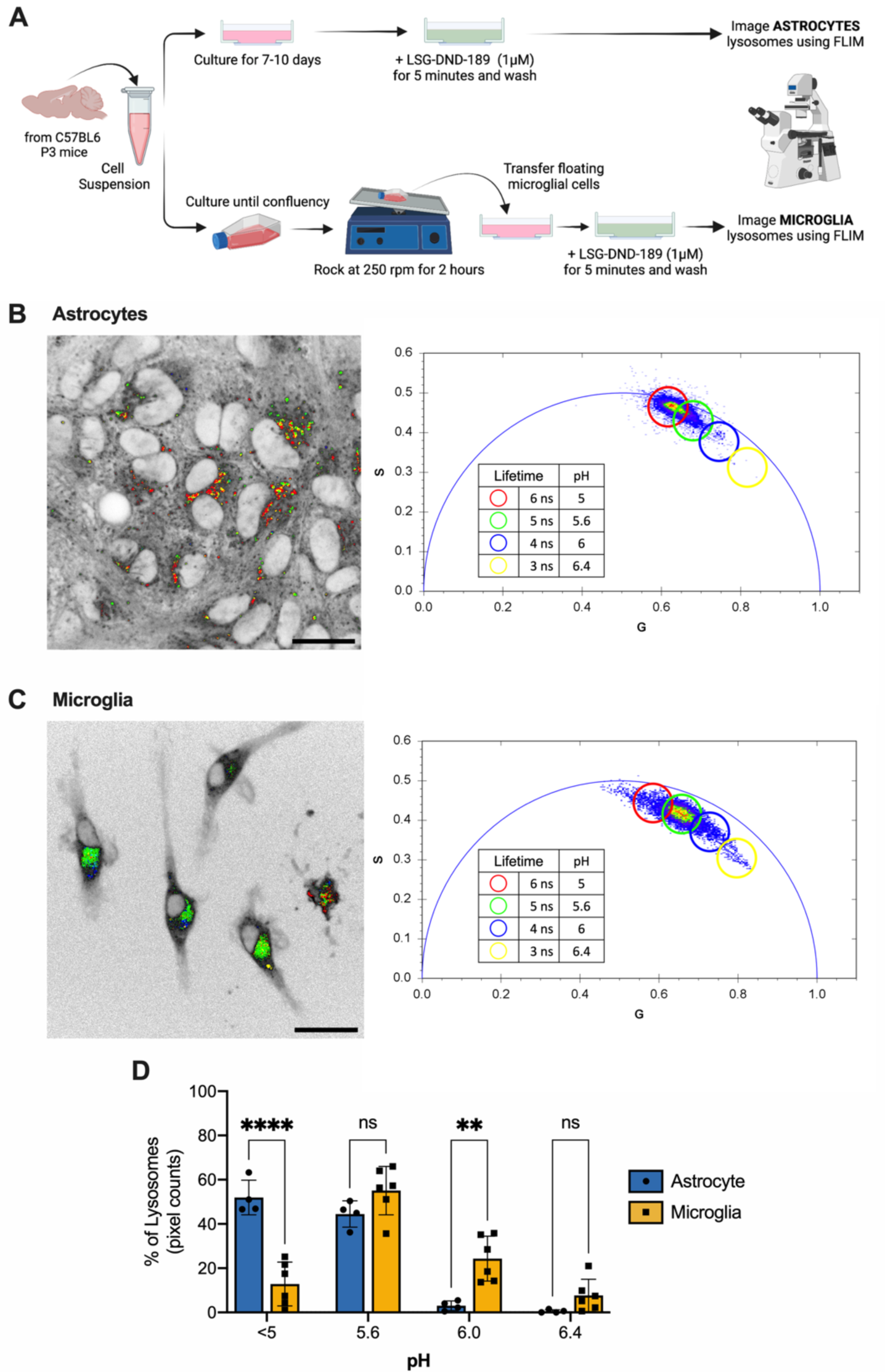
Lysosomal pH quantification in primary astrocytes and microglia. (A) Steps for preparation of primary astrocytes and microglia for LSG-DND-189 staining and FLIM imaging. (B and C) FLIM image and phasor plot for primary astrocytes and microglia. Pixels with different lifetime are pseudocolored in red (lifetime 6 ns and pH 5), green (lifetime 5 ns and pH 5.6), blue (lifetime 4 ns and pH 6) and yellow (lifetime 3 ns and pH 6.4). Scale bar, 20 μm. (D) Distribution of lysosomes across different pH in primary astrocytes and microglia. Data are mean ± SD, *\*\*\*\* p<0.0001*, ** *p<0.01, n.s.* not significant by two-way ANOVA with Sidak’s multiple comparison test.

We found that under control conditions, two types of glial cells, astrocytes and microglia show difference in lysosomal pH. First, when comparing the phasor plots of astrocytes and microglia, we noticed that the lifetime readings from microglial lysosomes span a larger area than those from astrocyte lysosomes (Figure 2, B and C). Upon pseudo-coloring of pixels with different lifetime/pH in FLIM images and grouping them in phasor plots, we then observed that astrocytes have more pixels in red that marks lysosomes with lifetime 6.0 ns and pH 5.0 (Figure 2B), while microglia showed more pixels pseudo-colored in green that marks lysosomes with lifetime 5.0 ns and pH 5.6 (Figure 2C). Next, quantitative analyses confirmed the difference in lysosomal pH distribution between astrocytes and microglia (Figure 2D). About 96% of astrocyte lysosomes are at pH 5.6 and lower, whereas 87% of microglial lysosomes are at pH 5.6 and higher (Figure 2D). Based on FLIM imaging and analyses, these results uncover the difference in lysosomal pH between two distinct glial cell types and suggest that lysosomal pH distribution provides an effective way in revealing lysosomal heterogeneity and assessing alterations in lysosomes.

### Monitoring lysosomal acidity dynamics in microglia upon different stimuli

External cues are known to play a crucial role in regulating lysosomal functions (Settembre and Ballabio, 2014). Deprivation of amino acids and growth factors induces autophagy, leads to the lysosomal activation of mTORC1, and subsequently results in the dilution of protons and increased lysosomal pH (Maulucci *et al*., 2015; Kwon *et al*., 2018, 2019). However, whether such deprivation affects lysosomal pH in primary glial cells remains unknown. We thus incubated primary microglia in Earl’s Balanced Salt Solution (EBSS), a nutrient-free medium that lacks amino acids and other growth factors, for 4 hours. Cells incubated in Dulbecco’s modified Eagle’s medium (DMEM) with complete nutrients serve as controls (Figure 3A). We found that upon starvation, the number of pixels pseudo-colored in blue that marks lysosomes with lifetime 4.0 ns and pH 6.0 increases, while those pseudo-colored in green representing lysosomes with lifetime 5.0 ns and pH 5.6 decrease their number (Figure 3B). Consistently, the percentage of lysosomes at pH 5.6 and pH 6.0 decreases and increases, respectively in starved microglial cells (Figure 3C).

**Figure 3.**
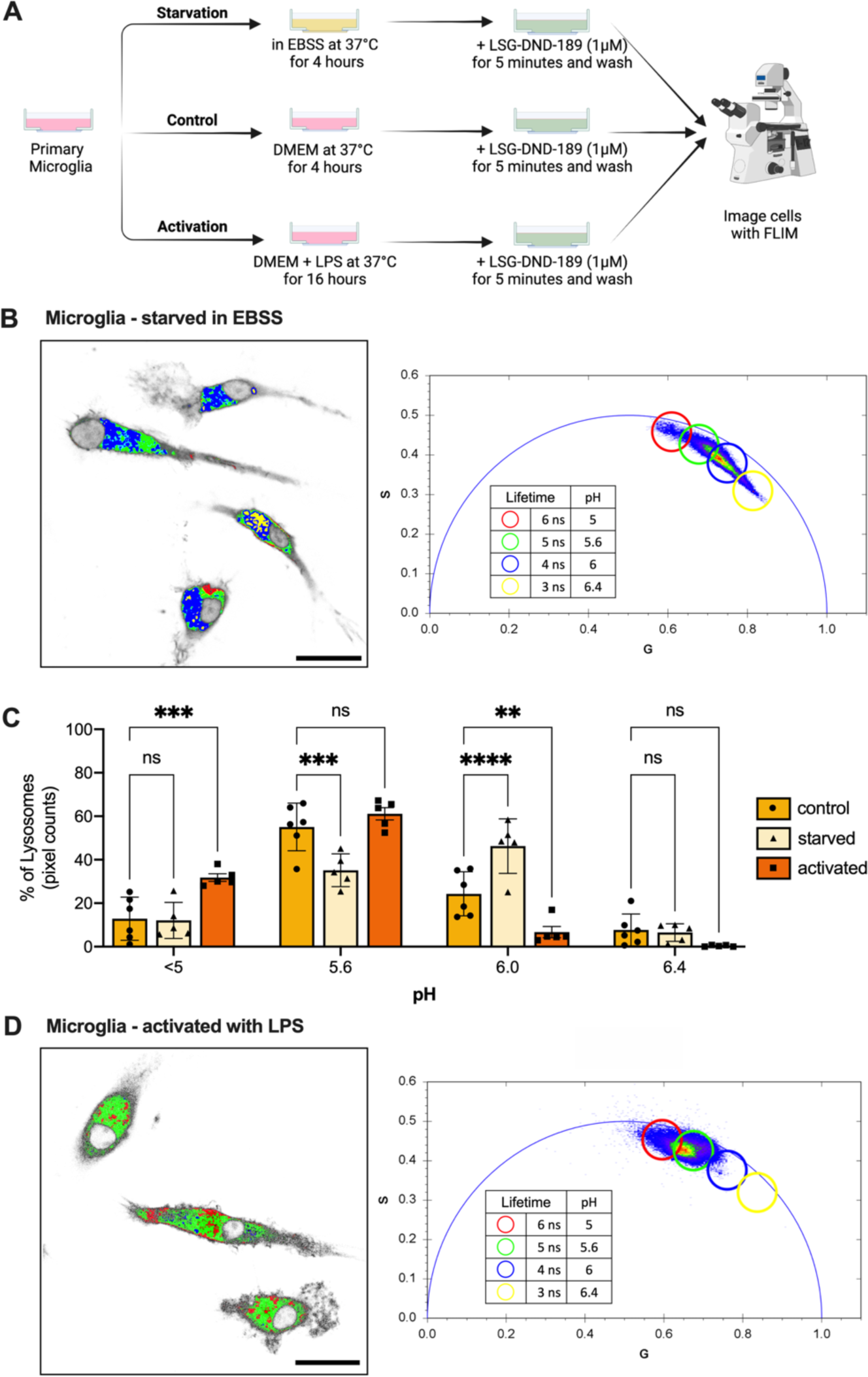
Lysosomal pH changes in primary microglia upon starvation and LPS activation. (A) Steps for preparation of primary microglia, including their starvation and activation by LPS, for LSG-DND-189 staining and FLIM imaging. (B) FLIM image and phasor plot for starved primary microglia. (C) Distribution of lysosomes across different pH in control, starved and activated microglia. Data are mean ± SD, *\*\*\*\* p<0.0001*, *\*\*\* p<0.001,* ** *p<0.01, n.s.* not significant by two-way ANOVA with Dunnett’s multiple comparison test. (D) FLIM image and phasor plot for primary microglia activated by LPS. For B and D, pixels with different lifetime are pseudocolored in red (lifetime 6ns and pH 5), green (lifetime 5 ns and pH 5.6), blue (lifetime 4 ns and pH 6) and yellow (lifetime 3 ns and pH 6.4). Scale bar, 20 μm.

On the other hand, astrocytes that were starved under similar conditions as microglia did not exhibit obvious lysosomal pH changes (Supplementary Figure 2, A-C). Although we observed a decrease in the number of lysosomes with pH 5.6 and pH 5.0 and an increase in lysosomes with pH 6.0, these changes were not statistically significant (*p>0.05).* Previous studies show that four hours of starvation in EBSS is sufficient to induce autophagy in primary astrocytes (Kulkarni *et al*., 2020). Therefore, the duration of starvation is unlikely to be the reason why we did not observe changes in lysosomal pH in primary astrocytes. These results thus suggest that microglia, as the major macrophages of the nervous system, may be more sensitive to nutritional stress in the environment, than astrocytes that are considered as supporting glial cells.

In addition, it is known that microglia can be activated by supplying lipopolysaccharide (LPS) into the culture medium (Herber *et al*., 2004; Majumdar *et al*., 2007). We observed that activated microglia have more red pseudo-colored pixels compared to control microglia, which suggests that LPS activation reduces lysosomal pH (Figure 3D). Further analysis quantified the numbers of pixels with different lifetime and revealed that LPS activation leads to a more than two-fold increase in the percentage of lysosomes at pH smaller than 5.0 and a more than two-fold decrease in those at pH 6.0 (Figure 3C). Our studies support the previous finding that the LPS treatment increases the acidity of microglial lysosomes (Majumdar *et al*., 2007) and further provide quantitative assessments regarding this crucial response. Together, these results not only demonstrate that microglial lysosomes accommodate with environmental changes by adjusting their acidity but also present the power of the FLIM imaging method in analyzing both decrease and increase of lysosomal pH. It would be interesting for future studies using this method to monitor how microglial lysosomes respond to other environmental stimuli and search for external cues that influence lysosomal pH in astrocytes.

### Neuron maturation associated with lysosomal acidification

Lysosomal activity and lysosome-related autophagy are required for neurogenesis, neuronal development and establishing neuronal polarity (Fassio *et al*., 2020). Neuronal lysosomes ensure proper membrane trafficking and axonal guidance, as well as synaptic functions (Lie and Nixon, 2018). To examine lysosomal acidity in neurons, we have isolated the brain from newborn P0 mice, dissociated neurons and started a primary neuron cell culture. During the process of neuron maturation, we have analyzed lysosomal pH using LSG-DND-189 labeling and FLIM imaging at day 6 and day 12 of culture time (Figure 4A).

**Figure 4.**
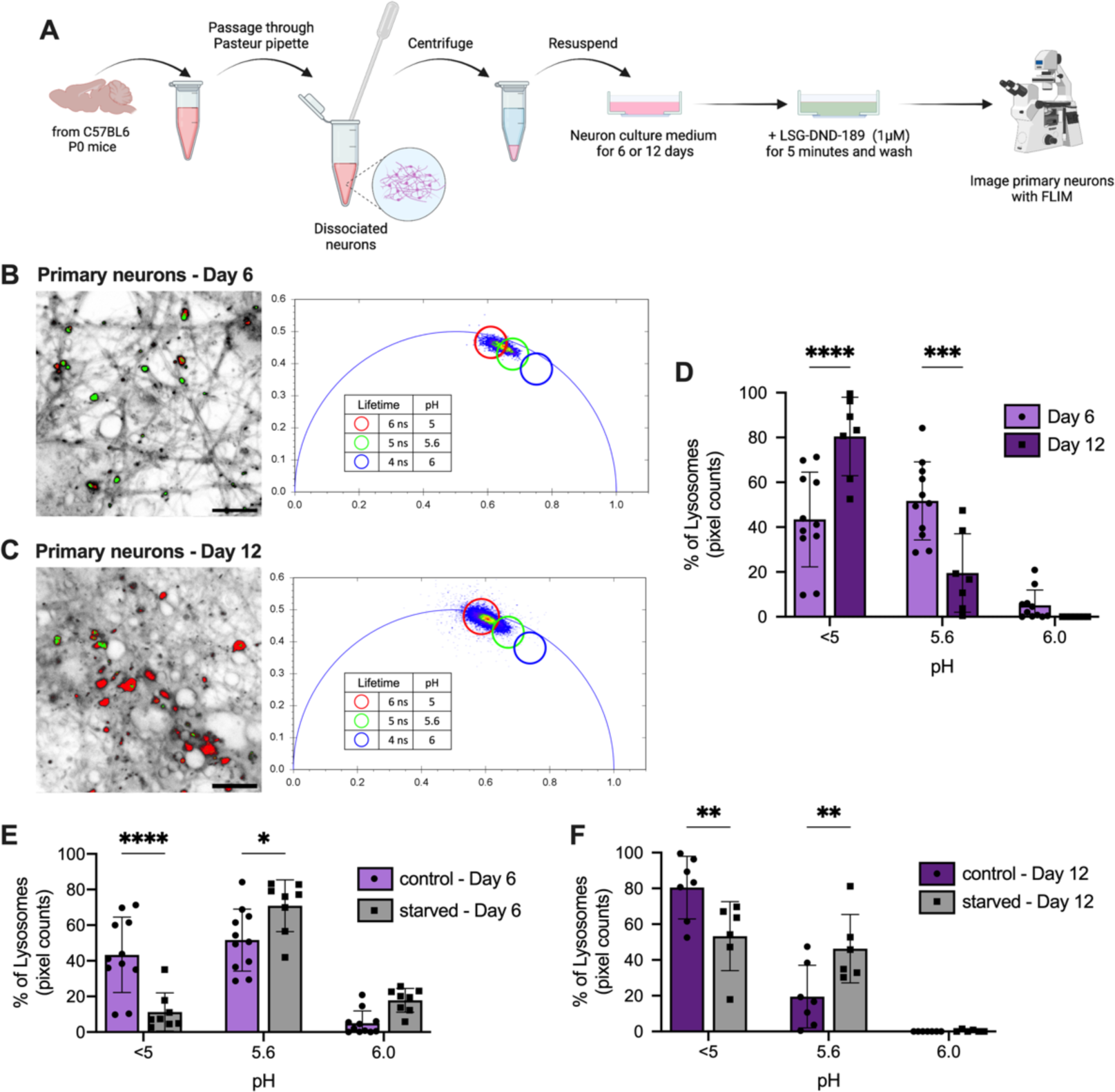
Lysosomal pH dynamics associated with primary neuron maturation and starvation. (A) Steps for preparation of primary neurons for LSG-DND-189 staining and FLIM imaging. (B and C) FLIM image and phasor plot of primary neurons at day 6 and day 12 of culturing. Pixels with different lifetime are pseudocolored in red (lifetime 6 ns and pH 5), green (lifetime 5 ns and pH 5.6) and blue (lifetime 4 ns and pH 6). Scale bar, 20 μm. (D) Distribution of lysosomes across different pH in day 6 and day 12 primary neurons. (E and F) Distribution of lysosomes across different pH upon starvation in day 6 and day 12 primary neurons. For D, E and F, data are mean ± SD, *\*\*\*\* p <0.0001*, *\*\*\* p <0.001,* ** *p < 0.01* by two-way ANOVA with Sidak’s multiple comparison test.

First thing we noticed is that the FLIM image and phasor plot of primary neurons are different than glial cells (Figure 2, B-C and Figure 4, B-C). The pixels on the phasor plot from neurons were more densely populated in a smaller region than those from glial cells. Most of these pixels from neurons had lifetimes >=6.0 ns and 5.0 ns indicating pH <5.0 and 5.6 respectively, and there were very few pixels with lifetime 4.0 ns indicating pH 6.0. Unlike glial cells, neurons showed no pixels with lifetime 3.0 ns indicating pH 6.4 (Figure 4D). Lysosomal pH distribution indicated clear differences among primary astrocytes, microglia, and neurons (Supplementary Figure 3A). Together, these results suggest that primary neurons have more homogenous and mature acidic lysosomes than glial cells. This observation is consistent with the notion that glial cells have more active endolysosomal pathways to provide nutritional support to neurons.

Furthermore, we found that lysosomes become more acidic as neurons mature from day 6 to day 12 in the culture. Day-6 neurons show more green pseudo-colored pixels with a lifetime of 5.0 ns indicating pH 5.6, whereas in day-12 neurons, pixels shift towards a longer lifetime of 6.0 ns indicating pH <5.0 (Figure 4, B and C). Almost 80% of the pixels in day-12 neurons have an average lifetime of 6.0 ns indicating pH <5.0 (Figure 4D). Thus, maturation of primary neurons leads to a decrease in lysosomal pH. Interestingly, previous studies showed that inhibition of lysosomal acidification in primary neurons triggers iron deficiency and causes neuronal stress by impairing mitochondrial function and promoting inflammation (Yambire *et al*., 2019). Therefore, we expect that increased acidity in mature neurons is required for their functionality, which may attribute to enhanced lysosomal degradation of macromolecules for the synthesis, release and recycle of signaling molecules involved in synaptic transmission and neural circuit formation.

Next, we asked whether starvation would also decrease lysosomal pH in primary neurons, as it does in microglia. We have used both day-6 and day-12 neurons and treated them in the EBSS medium for 4 hours. In the FLIM image and phasor plot, we observed a shift toward a shorter lifetime upon starvation, suggesting an increase in lysosomal pH. In both starved day-6 and day-12 neurons, there were more green pseudo-colored pixels with a lifetime of 5.0 ns/pH 5.6, but less red pseudo-colored pixels with a lifetime of 6.0 ns/pH<5.0 (Supplementary Figure 3, B and C, Figure 4, E and F). In day-6 neurons, the percentage of lysosomes with pH<5.0 was decreased to less than 10% upon starvation, while the percentage of lysosomes with pH 6.0 was increased to 20% (Figure 4E). However, in day-12 neurons, the percentage of lysosomes with pH<5.0 was maintained at more than 50%, and there were no lysosomes with pH 6.0 (Figure 4F). Together, these results suggest that as neurons become mature, their lysosomes increase acidity and exhibit increased resistance to amino acid and growth factor deprivation.

The starvation-induced increase in lysosomal pH has been observed in both microglia and neurons. Understanding the mechanism by which this increase occurs, and how this increase impacts lysosomal hydrolase activity and/or lysosomal signaling, ultimately leading to changes in microglial and neuronal functions, poses interesting questions for future studies.

### Tissue-specific variation in C. elegans lysosomal pH

In parallel with cellular imaging, we applied FLIM to analyze lysosomal pH in live organisms. To this end, we utilized *C. elegans*, a model organism with whole-body transparency, to facilitate direct visualization of lysosomes and their dynamics. We focused on two tissues, the intestine and hypodermis, in which lysosomes are accessible for LSG-DND-189 labeling and have been linked with metabolic and longevity regulations (McGhee, 2007; Folick *et al*., 2015; Baxi *et al*., 2017; Savini *et al*., 2019, 2022; Miao *et al*., 2020; Sun *et al*., 2020). Using FLIM, we imaged 1-day-old adult worms grown on LSG-DND-189 supplemented bacterial plates (Figure 5A). In the FLIM image and phasor plot, we observed a wide range of lifetime, varying from 1.5 ns to 5.0 ns. We found that the intestine contains more blue and purple pseudo-colored pixels with a lifetime of 3.0 ns/pH 6.4 and 2.0 ns/pH 6.8, respectively (Figure 5B). In contrast, all the red pseudo-colored pixels with lifetime 5.0 ns/pH 5.6 were localized in the hypodermis (Figure 5B). Further analyses showed that lysosomes in the intestine have a higher pH than those in the hypodermis (Figure 5C), revealing the diversity in lysosomal pH between these two tissues.

**Figure 5.**
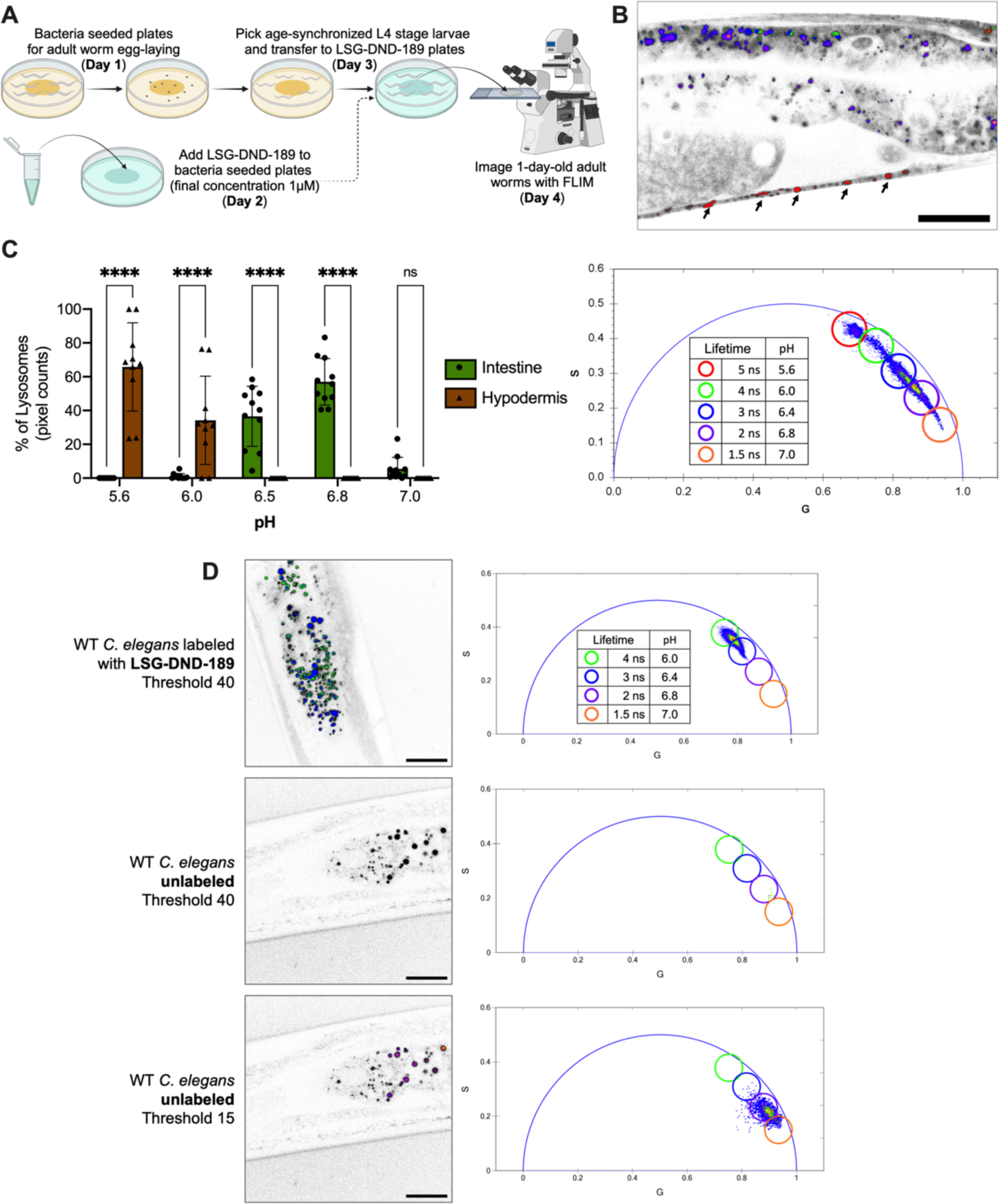
Lysosomal pH profiling in different tissues of *C. elegans*. (A) Steps to prepare *C. elegans* for LSG-DND-189 supplementation and FLIM imaging. (B) FLIM image and phasor plot of *C. elegans* supplied with LSG-DND-189. Arrows point to hypodermal lysosomes. (C) Distribution of lysosomes across different pH in *C. elegans* intestine and hypodermis. Data are mean ± SD, *\*\*\*\* p<0.001* and *n.s.* not significant by two-way ANOVA with Sidak’s multiple comparison test. (D) FLIM images and lifetime phasor plots of LSG-DND-189 labeled and unlabeled *C. elegans*. The FLIM images and lifetime phasor plots of unlabeled *C. elegans* are shown with two different thresholds, 40 and 15. FLIM images were acquired using the same laser intensity. For B and D, pixels with different lifetime are pseudocolored in red (lifetime 5 ns and pH 5.6), green (lifetime 4 ns and pH 6), blue (lifetime 3 ns and pH 6.4), purple (lifetime 2 ns and pH 6.8) and orange (lifetime 1.5 ns and pH 7.0). Scale bar, 20 μm.

These findings may imply that the intestine does not carry very acidic organelles in general. However, the *C. elegans* intestine is known to contain autofluorescent pigments, often referred to as lipofuscin or age pigment, which locate in gut granules (lysosome-like organelles). These gut granules can be stained with LysoTracker dyes and are dependent on lysosomal biogenesis genes for their formation (Clokey and Jacobson, 1986; Hermann *et al*., 2005). To investigate whether the autofluorescence originating from gut granules would affect the lifetime analysis, we imaged worms without LSG-DND-189 labeling. Although the photon counts of these autofluorescence signals were low, they exhibited a low lifetime of 1.5 ns, similar to the lifetime of the LSG-DND-189 signal at pH 7 (Figure 5D). Thus, there is a possibility that the high lysosomal pH observed in the *C. elegans* intestine is resulted from the presence of autofluorescence.

To further assess the contribution of autofluorescence and LSG-DND-189 to the combined lifetime signal in the intestine, we conducted a two-component analysis using phasor plots (Ranjit *et al*., 2019). We compared the phasor plots of LSG-DND-189 in pH buffer (Supplementary Figure 4A), in the intestine and hypodermis (Supplementary Figure 4B), as well as of autofluorescent gut granules in the unlabeled intestine (Supplementary Figure 4C). We found that the phasor cloud of the autofluorescent signal aligns with the line connecting the two phasor positions determined by measuring the lifetime of LSG-DND-189 within the pH buffer. The lifetime of the LSG-DND-189 signal from the intestine and hypodermis also corresponds to this same pattern. These results suggest that the elevated lysosomal pH observed in the intestine may arise from a complex interplay between autofluorescent pigments and LSG-DND-189. It is possible that the autofluorescence intensity could be enhanced upon LSG-DND-189 supplementation. Conversely, these autofluorescent pigments might influence the chemical properties of LSG-DND-189 within the intestine, potentially leading to a shift in its lifetime. Future experiments employing different lysosome pH-sensitive probes will be helpful in dissecting the impact of intestinal autofluorescence on lysosome pH measurements. At the current stage, the FLIM analysis in *C. elegans* is sufficient to determine changes in lysosomal pH but may not provide an accurate assessment of the absolute pH level.

Overall, our work demonstrates the effectiveness of FLIM in revealing lysosomal heterogeneity across various cell types and in live organisms. This FLIM-based method can measure small changes in lysosomal pH in both increasing and decreasing directions. Lysosomes house a variety of hydrolases that function optimally within a narrow pH range. Even subtle alterations in lysosomal pH can significantly impact the activity of these hydrolases. Through our analysis, we found that the distribution of lysosomal pH across the range is an effective and sensitive metric in revealing lysosomal heterogeneity and assessing subtle changes. The FLIM-based method is independent of fluorophore concentration, eliminating the need for a second reference fluorophore and ratiometric measurements. It also circumvents the requirement for transgenic reporters, although it does rely on the use of pH-sensitive probes that must be taken up by the cells, tissues, or organisms. Currently, the FLIM method using the LSG-DND-189 dye is the most sensitive for pH measurement in the range of 5.0 to 7.0, but it cannot further distinguish lysosomes with a pH below 5. Another LysoSensor dye DND-160, which has a pKa of 4.2, may work in the pH<5 range but requires excitation in the UV range, hindering its application in live cell imaging. Future applications of two-photon FLIM (2P-FLIM) technology in imaging the DND-160 dye, following a similar principle to what we demonstrate in this manuscript, may expand lysosomal pH measurements into a more acidic range. In parallel, the development of new lysosomal dyes with a pKa of 4.0 and a lifetime range distinct from the lifetime of autofluorescence pigments will be beneficial for FLIM imaging of lysosome pH in the *C. elegans* intestine and other mammalian tissues.

## MATERIALS AND METHODS

### FLIM instrumental set-up

The FLIM system employed in this study consists of an ISS Q2 laser scanning confocal nanoscope, one-photon excitation laser launcher, fastFLIM data acquisition and processing units. Prior to obtaining stable lifetime phasor plot measurements, the system requires a warm-up period of approximately 30 minutes. The fastFLIM technique operates in the frequency domain using Fourier transform analysis, therefore the initial phase and modulation of the FLIM system should be calibrated at specific modulation frequency. For calibration purposes in our study, we used a standard sample, Rhodamine 110, which has a lifetime of 4 ns. Rhodamine 110 was chosen due to the dynamic range of the LSG-DND-189 lifetime, which spans from approximately 1.5 ns to 6 ns. After calibration, the samples were imaged for lifetime measurement in frequency domain. Throughout our study, we used a 476 nm laser for excitation and a 500-633 nm emission filter for detection. The laser modulation frequency was set at 20 MHz. Imaging was conducted using a 60X water objective (Olympus, NA=1.20) with a field of view measuring 100 µm × 100 µm. The image resolution was configured to 1024 x 1024 pixels, with a pixel dwell time of 20 µs.

### Image acquisition and processing

The FLIM images were acquired and processed using the ISS VistaVision software and ImageJ. A flowchart illustrating the image processing steps can be found in Figure 1B. The FLIM image was processed using the phasor plot method in the frequency domain. Each pixel in the image with a specific g(ω) and s(ω) corresponds to a point in the phasor plot. A threshold was selected to eliminate the background signals. The background signal exhibits a Gaussian distribution; therefore, the higher edge of the Gaussian shape is chosen as the threshold to remove most of the background signals/pixels. Subsequently, pseudo-colors were assigned to highlight the distribution of lifetime for different populations. Although there may be a small overlap of pixels between these populations, the contribution of these pixels is considered negligible when compared to overall highlighted pixels. In the next step, all the pseudo-colored pixels associated with different lifetime/pH populations were merged to visualize the distributions of lysosomal pH. Finally, the pseudo-colored pixels with different lifetime/pH were quantified to determine the percentage of lysosomes within each pH range, thus generating a pH histogram.

### Primary astrocyte and microglia cell culture preparation and imaging

Primary glia cultures were prepared as described previously (Lian *et al*., 2016). Briefly, the cerebral cortices were isolated from P3 newborn pups (C57BL6 mice) in ice-cold dissection medium [Hanks’ balanced salt solution (HBSS) with 10 mM HEPES, 0.6% glucose, and 1% (v/v) penicillin/streptomycin], with meninges removed. The tissue was then finely minced and digested in 0.125% trypsin at 37°C for 15 minutes, followed by the addition of trypsin inhibitor (40 μg/mL) and DNase (250 μg/mL). Next, tissue was triturated, and resuspended in Dulbecco’s modified Eagle’s medium (DMEM) with 10% fetal bovine serum (FBS). The cell suspension was centrifuged and resuspended one more time to remove tissue debris. Cells were plated onto 35 mm glass bottom dishes (MatTek) pre-coated with poly-D-lysine (PDL) at a density of 50,000 cells/cm^2^ and cultured in DMEM with 10% FBS at 37°C in a humidified atmosphere of 95% air and 5% CO^2^ for 7-10 days. Media was changed every 3 days to suppress growth of microglia.

For microglia cultures, suspended cells were plated on T-75 flasks at a density of 50,000 cells/cm^2^ to generate mixed glial cultures. After the mixed glial culture reached confluency, the flasks were tapped against table and rocked at 250 rpm for 2 hours at 37 °C to separate microglia cells. The floating cells in media containing microglia were collected, seeded at 50,000 cells/cm^2^ in PDL coated 35 mm glass bottom dishes and grown in DMEM with 10% FBS at 37°C for two more days before experiments. For LPS treatment, 200 ng/ml LPS (Sigma-Aldrich) was added to the microglia culture media 16 hours before an experiment.

On the day of imaging, the cells were stained with 1 μM LysoSensor Green DND-189 (Thermo Fisher) in media for 5 minutes. Cells were rinsed twice with 1X PBS and incubated in culture medium for FLIM measurement.

### Neuron cell culture preparation and imaging

Primary neuronal cultures were prepared from P0 mouse (C57BL6 mice). The cerebral cortices were dissected and minced in ice-cold dissection medium and treated with 0.125% trypsin at 37°C for 15 minutes as above. The reaction was stopped with trypsin inhibitor solution (40 μg/mL) containing DNAse (250 μg/mL). Neurons were dissociated by several passages through a Pasteur pipette. The cells were then centrifuged at 1000 x g for 5 minutes, resuspended in neuron culture media (Neurobasal medium supplemented with 2% B27, 0.5 mM L-glutamine, 0.4% v/v Pen/Strep). The cell suspension was centrifuged and resuspended one more time to remove tissue debris. The washed neurons were plated on 35mm glass bottom dishes precoated with PDL at a density of 50,000 cells/cm2 and incubated at 37 °C in a humidified atmosphere of 95% air and 5% CO2 for five or twelve days before experiments. On the day of imaging, the cells were stained with 1 μM LysoSensor Green DND-189 (Thermo Fisher) in media for 5 minutes. Cells were rinsed twice with 1X PBS and incubated in culture medium for FLIM measurement.

### Starvation assay in primary glia and neurons

On the day of imaging, for amino acid and serum starvation, cells were incubated in pre-warmed EBSS (Earls Balanced Salt Solution) (Invitrogen) at 37°C for 4 hours to induce autophagy. LysoSensor Green DND-189 stock solution (Thermo Fisher) was diluted to the final working concentration (1 μM) in either normal cell culture medium or EBSS. The cells were stained with 1 μM LysoSensor Green DND-189 in media for 5 minutes. Cells were rinsed twice with 1X PBS and incubated in culture medium for FLIM measurement.

### Preparation and starvation of C. elegans

*C. elegans* wild-type (N2) strain was maintained on standard nematode growth media (NGM) plates seeded with OP50 *E. coli* bacteria (at 20°C. LysoSensor Green DND-189 (Thermo Fisher) was diluted in M9 buffer and added to 6 cm standard NGM plates (containing 12 mL of agar) seeded with bacteria, to a final concentration of 1 mM. The plates were kept in the dark for 24 hours to allow the LysoSensor solution to diffuse evenly throughout the plate. Age-synchronized larval L4 stage *C. elegans* were added to these plates and maintained for 24 hours in the dark before imaging with FLIM. For imaging, worms were mounted on glass slides with 2% agarose pads containing 0.5% NaN^3^ as anesthetic.

### Quantification and statistical analysis

No statistical method was used to pre-determine sample size. Cells and animals were randomly picked from experimental groups for analysis, and data from all the samples were used for statistics without exclusion. Investigators were not blinded. Student’s t-test was used when comparing two samples. Two-way ANOVA was performed with Sidak’s multiple comparison test when two effectors, for example cell type and lysosomal pH distribution, were compared. Two-way ANOVA was performed with Dunnett’s test when multiple conditions were compared to a control, for example microglia under control, starvation, and activation conditions. All statistical analysis was done in PRISM 9 by GraphPad.

## ACKNOWLEDGEMENTS

We thank Princeton Svay for providing technical support. We extend our appreciation to Dr. Yong Yu and Guo Hu for their valuable experimental support. We also acknowledge the insightful discussions about the FLIM system with Drs. Yuansheng Sun and Shih-Chu Jeff Liao. This work was supported by NIH grants RF1AG074540 (A.S.M.), P01AG066606 (A.S.M. and H.Z.) and RF1NS093652 (H.Z). M.C.W was supported by the Janelia Research Campus, Howard Hughes Medical Institute.

## SUPPLEMENTARY FIGURES

**Supplementary Figure 1.**
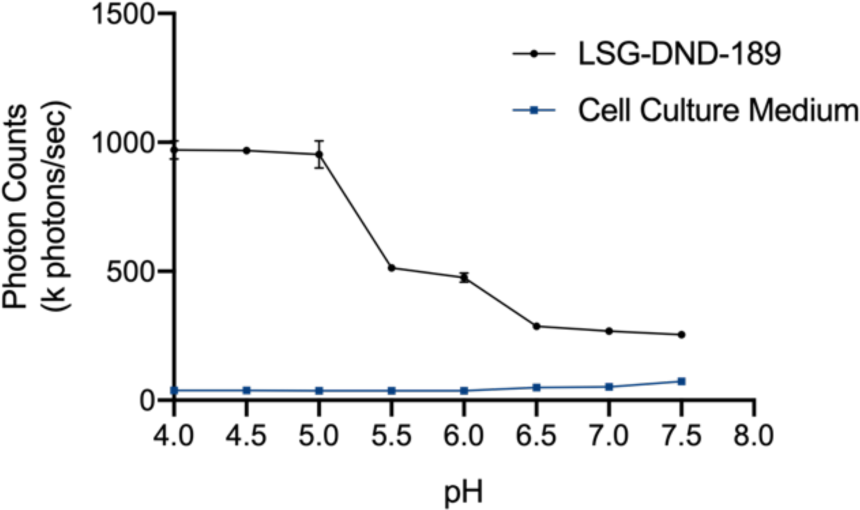
Average fluorescence intensity of LSG-DND-189 and cell culture medium across difference pH as represented by photon counts.

**Supplementary Figure 2.**
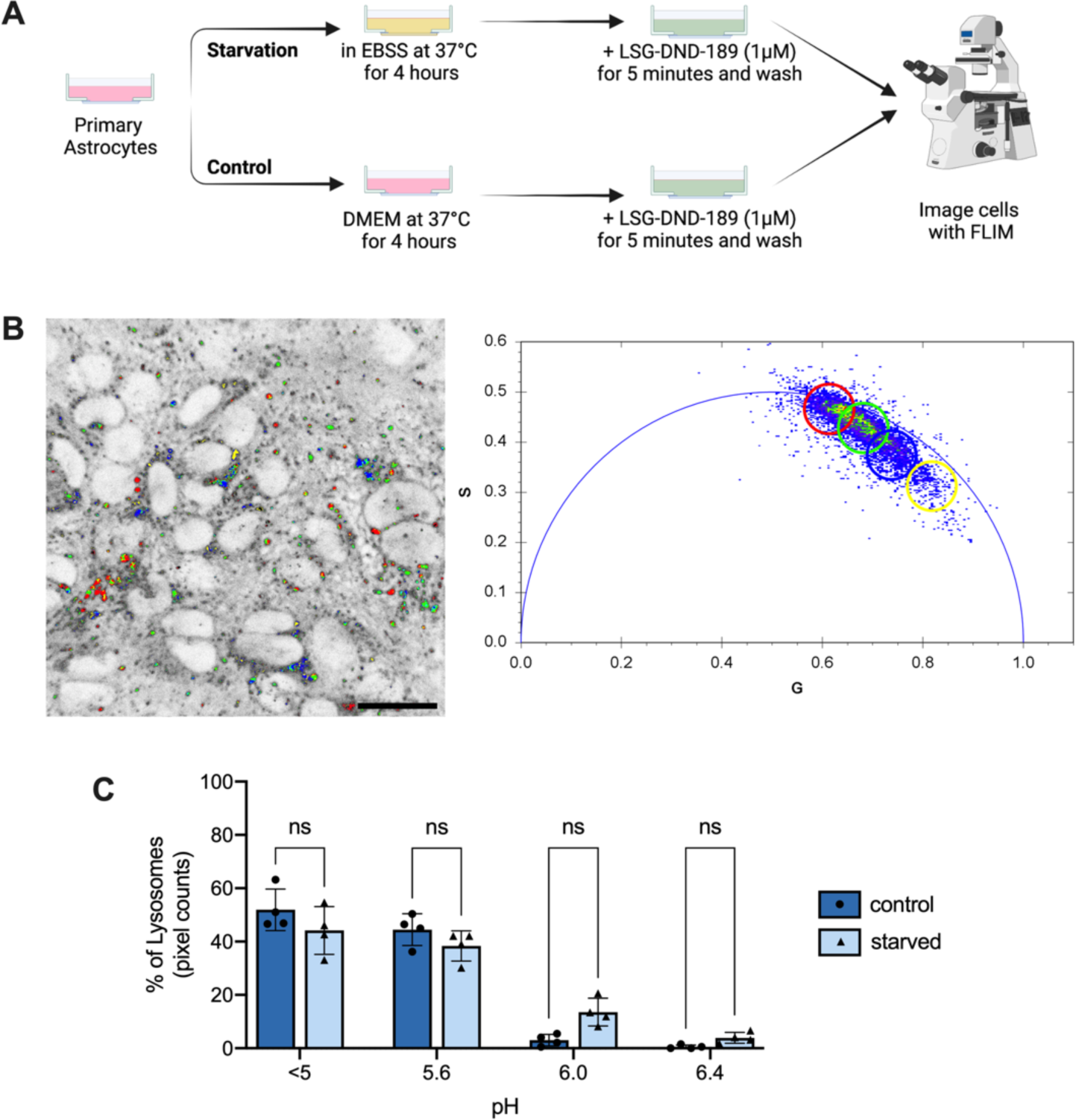
Lysosomal pH in primary astrocytes upon starvation. (A) Steps for preparation of primary astrocytes for starvation, LSG-DND-189 staining and FLIM imaging. (B) FLIM image and phasor plot of starved primary astrocytes. Pixels with different lifetime are pseudocolored in red (lifetime 6 ns and pH 5), green (lifetime 5 ns and pH 5.6), blue (lifetime 4 ns and pH 6) and yellow (lifetime 3 ns and pH 6.4). Scale bar, 20 μm. (C) Distribution of lysosomes across different pH in control and starved primary astrocytes. Data are mean ± SD, *n.s.* not significant by two-way ANOVA with Sidak’s multiple comparison test.

**Supplementary Figure 3.**
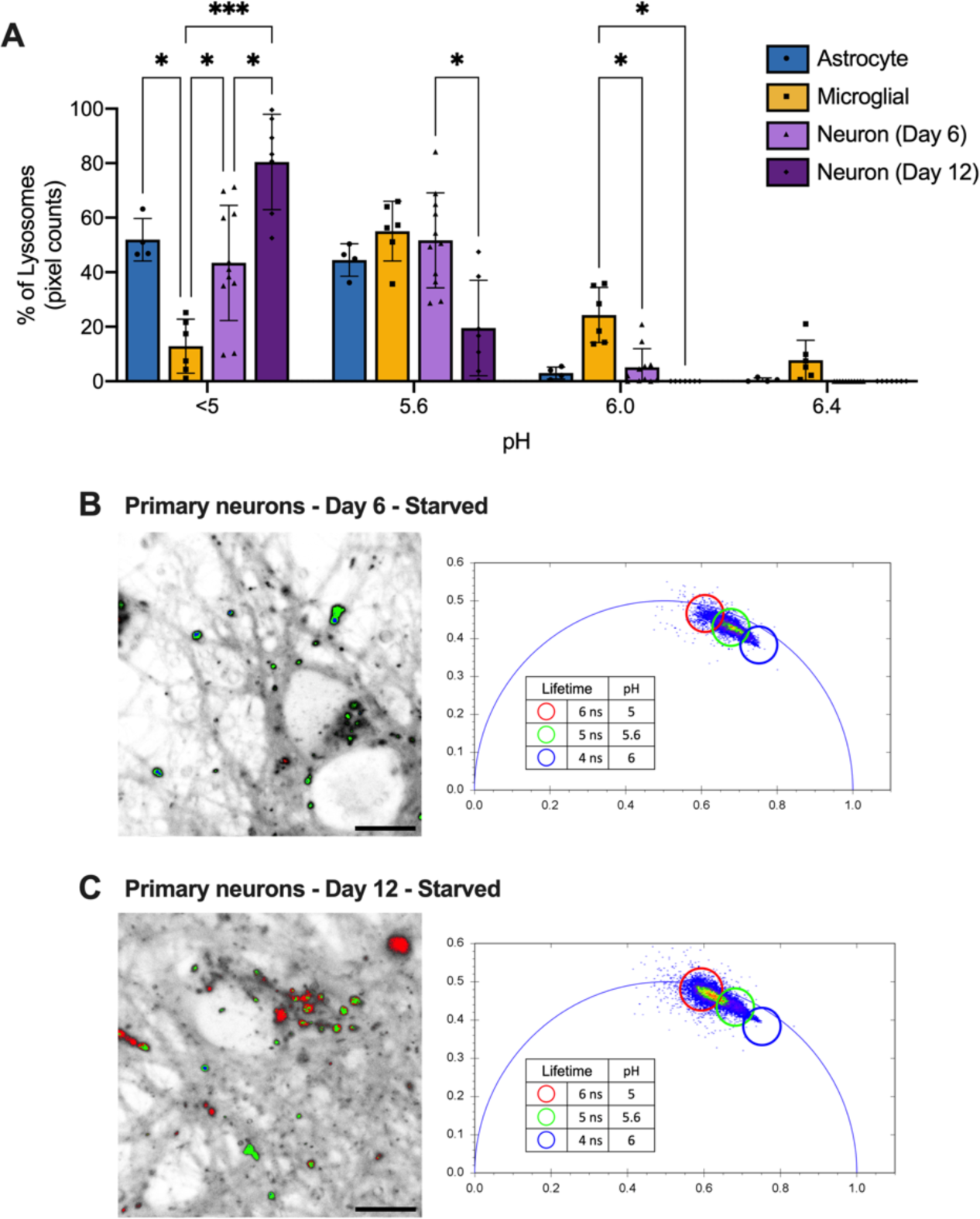
Lysosomal pH in starved primary neurons. (A) Distribution of lysosomes across different pH in primary astrocytes, microglia, and neurons. Data are mean ± SD, *\*\*\*\* p<0.0001, * p<0.05* by two-way ANOVA with Sidak’s multiple comparison test. (B and C) FLIM image and phasor plot of starved primary neurons in day 6 and day 12 of culturing. Pixels with different lifetime are pseudocolored in red (lifetime 6 ns and pH 5), green (lifetime 5 ns and pH 5.6) and blue (lifetime 4 ns and pH 6). Scale bar, 20 μm.

**Supplementary Figure 4.**
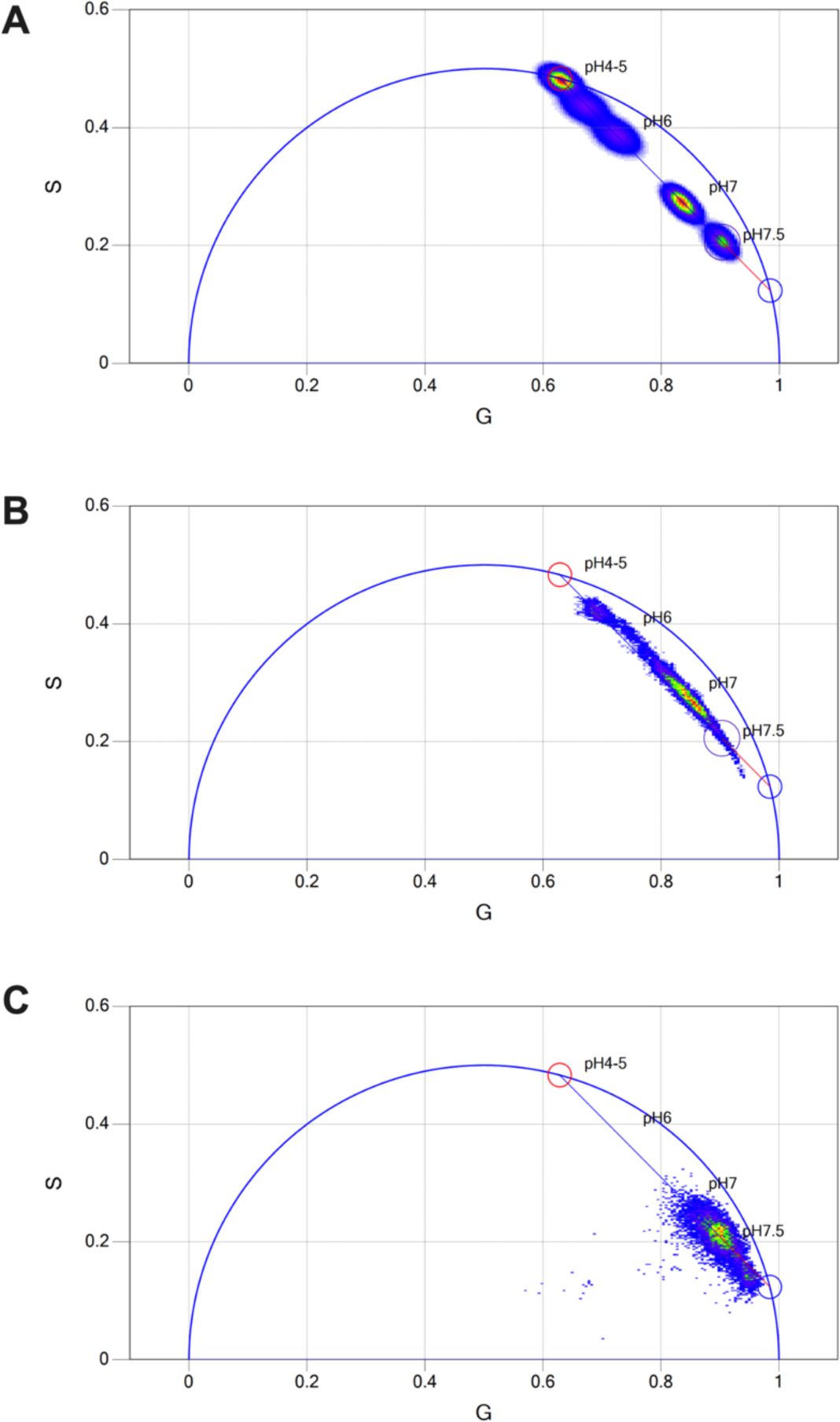
Two-component analysis using lifetime phasor plots of (A) LSG-DND-189 in pH buffer, (B) LSG-DND-189 in *C. elegans* and (C) autofluorescence in unlabeled *C. elegans*.

## Notes

### Competing Interest Statement

The authors have declared no competing interest.

